# A proximity-labeling proteomic approach to investigate invadopodia molecular landscape in breast cancer cells

**DOI:** 10.1101/869545

**Authors:** Sylvie Thuault, Claire Mamelonet, Joëlle Salameh, Kevin Ostacolo, Brice Chanez, Danièle Salaün, Emilie Baudelet, Stéphane Audebert, Luc Camoin, Ali Badache

## Abstract

Metastatic progression is the leading cause of mortality in breast cancer. Invasive tumor cells develop invadopodia to travel through basement membranes and the interstitial matrix. Substantial efforts have been made to characterize invadopodia molecular composition. However, their full molecular identity is still missing due to the difficulty in isolating them. To fill this gap, we developed a non-hypothesis driven proteomic approach based on the BioID proximity biotinylation technology, using the invadopodia-specific protein Tks5α fused to the promiscuous biotin ligase BirA* as bait. In invasive breast cancer cells, Tks5α fusion concentrated to invadopodia and selectively biotinylated invadopodia components, in contrast to a fusion which lacked the membrane-targeting PX domain (Tks5β). Biotinylated proteins were isolated by affinity capture and identified by mass spectrometry. We identified known invadopodia components, revealing the pertinence of our strategy. Furthermore, we observed that Tks5 newly identified close neighbors belonged to a biologically relevant network centered on actin cytoskeleton organization. Analysis of Tks5β interactome demonstrated that some partners bound Tks5 before its recruitment to invadopodia. Thus, the present strategy allowed us to identify novel Tks5 partners that were not identified by traditional approaches and could help get a more comprehensive picture of invadopodia molecular landscape.

## INTRODUCTION

While primary breast tumor patients can often be cured by surgery and/or radiation therapy combined to chemotherapy when detected at early stages, progression of the disease to the metastatic stage is still a major cause of patient death. Therefore, deciphering the molecular mechanisms underlying metastatic progression is of crucial importance for a better control of the disease. In order to reach the vascular system and be transported to distant organs and colonize them, tumor cells need to breach through basement membranes and invade through the interstitial matrix. Extracellular matrix (ECM) proteolysis is required to clear space and create tracks along which cells migrate. Invadopodia, which are dynamic actin-rich protrusive structures that cancer cells develop at their surface, play a major role in ECM proteolysis^1–3^. Furthermore, increasing amount of evidence point to their involvement in metastasis^4,5^. Therefore, a better understanding of their biology and molecular characteristics might permit the identification of markers of metastatic propensity and potential therapeutic targets.

Major efforts have been made during the past 25 years to characterize invadopodia molecular composition, using classical targeted proteomics involving cell fractionation or co-immunoprecipitation after cell lysis or candidate approaches. These studies have established that invadopodia are composed of structural and signaling proteins, including cortactin, cofilin, N-WASP and Tks4/5, that control the reorganization of the underlying actin cytoskeleton and the release of proteases such as MT1-MMP, involved in matrix degradation^1–3^. Furthermore, time-lapse microscopy analyses revealed that invadopodia formation and maturation involve three major steps: initiation, assembly and maturation^6,7^. However, the full molecular identity of these structures is missing due to the fact that their isolation and purification have been challenging^8^.

To circumvent these limits, we developed a non-hypothesis driven approach based on the proximity-dependent BioID biotinylation assay, coupled to mass spectrometry analysis^9^. This technique relies on the engineering of cells expressing a protein of interest fused to the promiscuous biotin ligase BirA*. In presence of biotin, BirA* biotinylates its close protein neighbors *in situ*, which can then be isolated using avidin-based affinity capture and identified by mass spectrometry. Proximity-dependent techniques allow to explore the molecular landscape of membrane-less organelles or of protein complexes localized to poorly soluble subcellular compartments, which are poorly accessible by more established approaches such as cell fractionation and affinity purification. Moreover, they enable the identification of weak or transient protein-protein interactions^10,11^.

We wished to establish whether proximity labelling could help characterize invadopodia at the molecular level. For that purpose, the scaffold protein Tks5, also known as SH3PXD2A or FISH, was used as bait. Indeed, in contrast to other invadopodia components which are also components of other actin-based structures such as lamellipodia or focal adhesions, Tks5 is specific of invadopodia. Tks5 protein contains five Src Homology 3 (SH3) domains, proline-rich regions (PxxP) and, depending on the isoform, a N-terminal phox-homology (PX) domain which confers affinity for phosphatidylinositol-3,4-bisphosphate (PI(3,4)P2)^12,13^. Three Tks5 isoforms have been reported arising from distinct promoter usage: Tks5α (also referred as Tks5long), Tks5β and Tks5short^14–16^. Tks5α is the only isoform containing the N-terminal PX domain. In cancer, the expression level of Tks5α or the ratio of Tks5α expression to Tks5β/Tks5short expression, rather than the total level of Tks5, is correlated to poor prognosis and lower metastasis-free survival in diverse cancer types^15,17–20^. This is correlated with the fact that Tks5α plays a major role in invadopodia formation and cancer cell invasion. Indeed, consequently to its recruitment at PI(3,4)P_2_-enriched membrane domains, Tks5 potentiates actin polymerization through its interaction with the WASP-Arp2/3 complex^21^. Moreover, Tks5α through direct or indirect recruitment of proteases at invadopodia contributes to ECM degradation^22,23^. Finally, Tks5 facilitates the local generation of reactive oxygen species which are required for podosome (counterpart of invadopodia in normal cells) and invadopodia formation^24^.

In this study, in order to determine Tks5 interactome *in situ* and therefore invadopodia molecular composition, we engineered MDA-MB-231 breast cancer cell lines expressing either Tks5α, or Tks5α lacking the PX domain fused to the biotin ligase BirA* and verified that this allowed the biotinylation and identification of invadopodia components. Label-free mass spectrometry analysis identified new potential invadopodia proteins that formed a protein-protein interaction network of actin cytoskeleton regulators. Tks5α lacking the PX domain recruited an overlapping but distinct group of proteins indicating that some proteins associate to Tks5 before its recruitment to invadosomes. Thus, the present strategy shows great potential for the molecular characterization of invadopodia *in situ*.

## RESULTS

### Selective biotinylation of invadopodia proteins by Tks5-BirA* fusion proteins

In order to characterize invadopodia at the molecular level, we set up the identification of the close neighbors of a specific component of invadopodia, the adaptor protein Tks5α^12^, using the BioID proximity ligation technology. We therefore stably expressed wild type Tks5α (Tks5) fused to BirA* in MDA-MB-231 breast cancer cells (Fig.1A). We also generated MDA-MB-231 cells expressing a mutant form of Tks5α lacking its PX domain (ΔPX-Tks5) fused to BirA* (Fig.1A). This mutant, mimicking the Tks5β/short form of Tks5, should not localize to invadopodia. Tks5β has also been described to have a dominant negative effect over Tks5α^15^. Finally, cells expressing BirA* alone were established as control for identification of unspecific biotinylation. We derived individual clones by limiting dilution and selected cells with similar expression levels of the fusion proteins (Fig.1C). Invasive cells form invadopodia when plated on an artificial ECM. Tks5-BirA* expressing cells were seeded on fluorescently-labeled gelatin to visualize invadopodia formation. We observed that Tks5-BirA* localized at mature invadopodia, identified by cortactin labeling and gelatin degradation, whereas ΔPX-Tks5-BirA* and BirA* did not (Fig.1B). Moreover, as previously reported, Tks5 overexpression increased the ability of cells to degrade gelatin^13,25^. Indeed, we observed an increase in the percentage of cells degrading the matrix, but also in the area that was degraded per individual cells (Fig.1B, D-E). In contrast, expression of ΔPX-Tks5 reduced the percentage of degrading cells. These observations demonstrate that Tks5-BirA* fusion proteins are functional and that only full-length Tks5-BirA* is strongly enriched in invadopodia.

**Figure 1:**
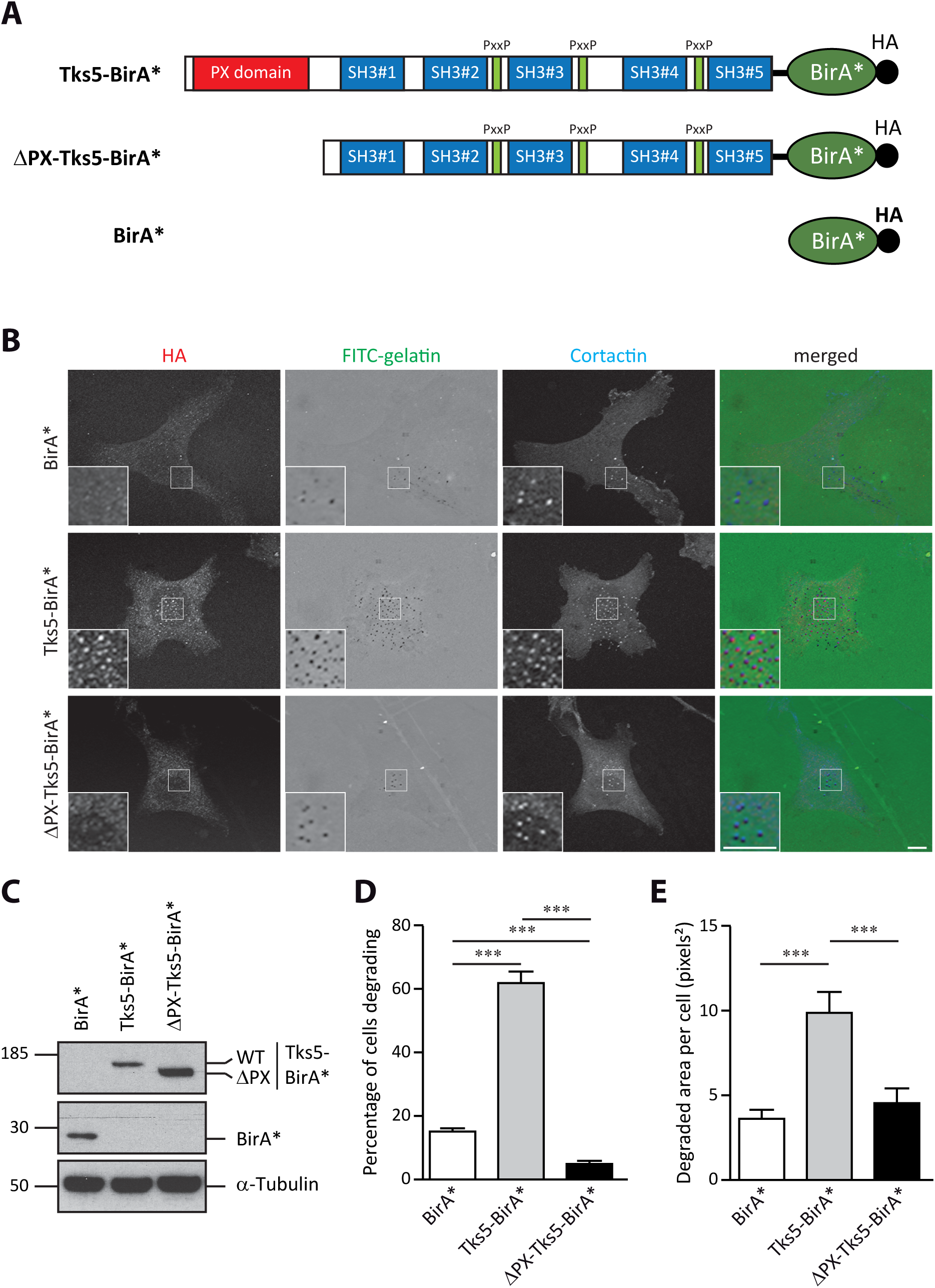
Tks5-BirA* is active and localizes in invadopodia. (A) Schematic representation of BirA* fusion proteins used in this study. The PX, SH3 and Proline rich (PxxP) domains of Tks5 are depicted. All fusion proteins have a HA tag at their C-terminus. (B) Representative images of MDA-MB-231 cells stably expressing wild-type Tks5 or Tks5 lacking its PX domain (ΔPX) fused to the biotin ligase BirA* or BirA* alone, seeded on fluorescently-labeled gelatin (FITC-gelatin) for 4 hours. Cells were fixed and stained with anti-HA and -cortactin antibodies. Invadopodia were identified thanks to cortactin labeling. Active invadopodia were identified thanks to degradation area (dark spots) in FITC-gelatin. The white-boxed regions are shown enlarged in the bottom insets. Scale bars represent 5μm. (C) Expression levels of BirA* fusion proteins in MDA-MB-231-derived cells described in (B) was assessed by western blotting using an anti-HA antibody. α-Tubulin was used as a loading control. (D-E) Tks5-BirA* fusion proteins are functional. The ability of MDA-MB-231 cells expressing Tks5-BirA* fusion proteins, described in (B), to degrade fluorescently-labeled gelatin was analyzed. The percentage of degrading cells (D) and the degraded area per cell (E) are represented as the mean ± SEM of three independent experiments. *** p≤0.001.

It was then important to verify that Tks5-BirA* could induce selective biotinylation at invadopodia. Cells were seeded on fluorescent gelatin in presence of biotin for 16 hours and localization of biotinylated proteins was assessed by staining with streptavidin coupled to Alexa Fluor® 594. We observed an accumulation of biotinylated proteins at invadopodia in cells expressing Tks5-BirA*, but not in cells expressing the mutant ΔPX-Tks5-BirA* or BirA* alone (Fig.2A and data not shown). In cells expressing ΔPX-Tks5-BirA*, biotinylated proteins were exclusively cytoplasmic. Thus this approach was likely to reveal invadopodia components.

**Figure 2:**
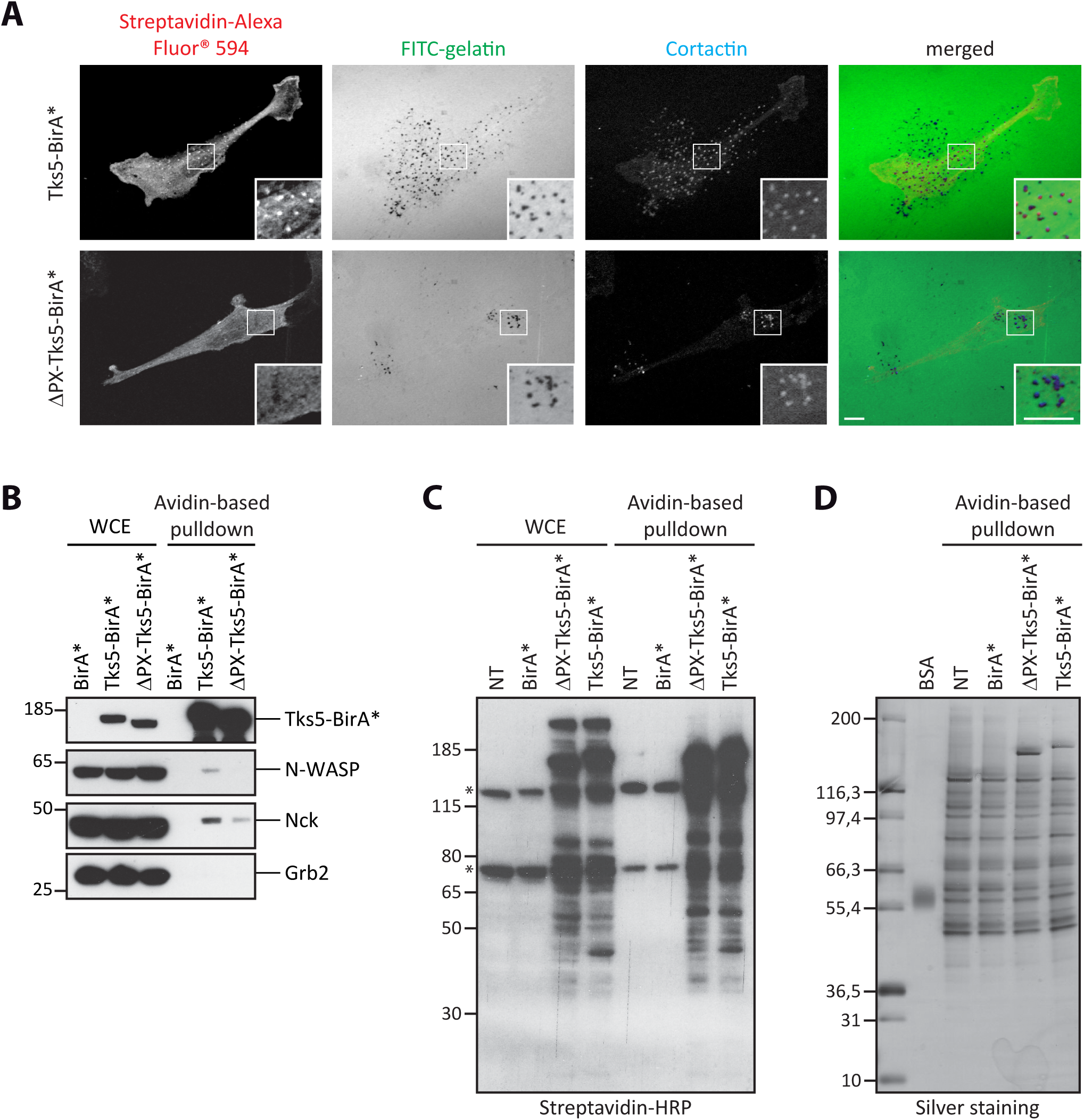
Selective biotinylation of invadopodia proteins by Tks5-BirA*. (A) Representative images of MDA-MB-231 cells stably expressing Tks5-BirA* or ΔPX-Tks5-BirA* seeded on fluorescently-labeled gelatin (FITC-gelatin) in the presence of biotin for 16 hours. Cells were fixed and stained for cortactin as a marker of invadopodia and streptavidin coupled to Alexa Fluor® 594 to reveal biotinylated proteins. The white-boxed regions are shown enlarged in the bottom insets. Scale bars represent 5μm. (B) Analysis of the biotinylation of known Tks5 interactors and of well characterized invadosome components. MDA-MB-231 cells stably expressing BirA*, Tks5-BirA* or ΔPX-Tks5-BirA* were seeded on gelatin in presence of biotin for 16 h. After cell lysis, biotinylated proteins were isolated by affinity capture using avidin-coated beads. The presence of proteins of interest was assessed in whole cell extract (WCE) or after avidin-based pulldown by western blotting with specific antibodies. (C) Isolation of proteins biotinylated by Tks5-BirA* and ΔPX-Tks5-BirA*. MDA-MB-231 cells stably expressing Tks5-BirA*, ΔPX-Tks5-BirA* or BirA*, and non-transfected (NT) cells were seeded on gelatin and treated with biotin for 16 h before cell lysis. Biotinylated proteins were pulled down via avidin-coated beads. The presence of biotinylated proteins was assessed by western blotting in whole cell extract (WCE) or after avidin-based pulldown (one-twentieth of the total pulldown was used) using HRP-coupled streptavidin. (*) corresponds to intrinsically biotinylated proteins. (D) One-tenth of the samples shown in (C) was resolved by SDS-PAGE for silver staining of the isolated proteins; the rest was analyzed by mass spectrometry.

Finally, we assessed if previously described partners of Tks5 and invadosomes components, such as N-WASP, Nck and Grb2^21,25^, could indeed be biotinylated by Tks5-BirA*. MDA-MB-231 cells expressing Tks5-BirA* fusion proteins or BirA* were seeded on gelatin for 16 hours in the presence of biotin. Biotinylated proteins were isolated by affinity capture using avidin-coated beads and the biotinylation of the proteins of interest assessed by western blotting. N-WASP and Nck were biotinylated by Tks5-BirA* but not, or much less, by ΔPX-Tks5-BirA* (Fig.2B). None of the proteins were biotinylated by BirA*. Thus our approach is capable of identifying *bona fide* Tks5 partners. This result also shows that the interactions of Tks5 with Nck and N-WASP, that have been previously identified by co-immunoprecipitation after cell lysis, also occurred *in vivo*. Furthermore, the fact that N-WASP and Nck were poorly biotinylated by ΔPX-Tks5-BirA* strongly suggested that their interaction with Tks5 occurred once Tks5 had been recruited at PI(3,4)P_2_-enriched invadopodia. We cannot totally exclude, however, that N-WASP and Nck require the presence of the PX domain for the interaction. By contrast, Grb2 was not biotinylated by Tks5-BirA* fusion proteins (Fig.2B). This observation is in agreement with a previous study indicating that Grb2 can distinguish between invadopodia and podosomes^26^. The lack of Grb2 biotinylation could also be due to improper relative orientation of the fused proteins and/or inaccessible lysine residues. Altogether these observations show that MDA-MB-231 cells expressing Tks5-BirA* and ΔPX-Tks5-BirA* are suitable cell models to address invadopodia composition in a global manner.

### Tks5α close neighbors belong to an integrative network associated with actin cytoskeleton reorganization

Analysis of biotinylation in whole cell extracts or after affinity capture indicated that Tks5-BirA* fusion proteins led to high levels of biotinylation of a restricted number of proteins, whereas few proteins were biotinylated to detection levels in non-transfected MDA-MB-231 cells or cells expressing BirA* alone (Fig.2C). Silver staining of the proteins pulled down in non-transfected cells or cells expressing BirA* revealed a substantial number of proteins binding non-specifically to avidin-coupled beads (Fig.2D). We next conducted mass spectrometry analysis of affinity captured proteins from Tks5-BirA* expressing cells and control cells. Relative intensity-based label-free quantification (LFQ) method identified 40 proteins significantly enriched with a 1% permutation FDR (pFDR) in Tks5-BirA* pulldown compared to BirA* control pulldown (Fig.3A and Supplementary Table S1).

**Figure 3:**
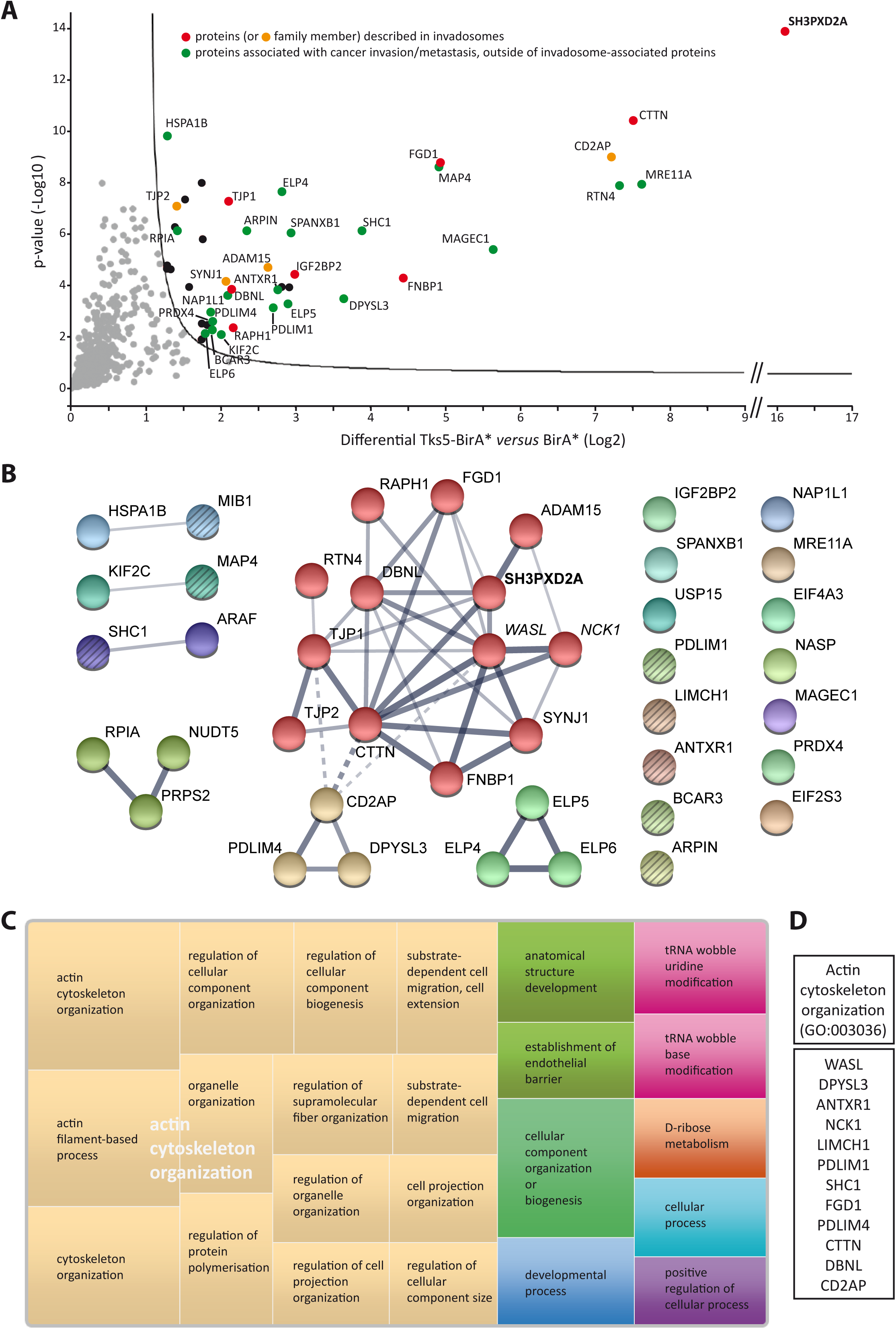
Tks5α close neighbors belong to an integrative network related to actin cytoskeleton reorganization. MDA-MB-231 cells expressing Tks5-BirA* or BirA* were seeded on gelatin-coated plates and treated with biotin for 16 h. Biotinylated proteins were isolated by affinity capture and identified by mass spectrometry. (A) Volcano plot showing the differential level (Log2) and the p-value (-Log10), on x and y axis, respectively, for Tks5-BirA* *versus* BirA*. The full line is indicative of protein hits obtained at a permutation false discovery rate (pFDR) of 1%. Data result from two different experiments processed three times. Red and orange spots highlight proteins previously associated with invadosomes and proteins for which a family member has been associated with invadosomes, respectively. Green spots highlight proteins associated with cancer invasion/metastasis, outside of invadosome-associated proteins. For more details, refer to Supplementary Table S1. (B) Proteins biotinylated by Tks5-BirA*, including the validated candidate proteins (WASL and NCK1 which appear in italic) were integrated in a protein-protein interaction network based on physical and functional interactions per the STRING database. Markov Clustering algorithm has been applied. Proteins belonging to a same cluster have an identical color; the main core cluster is in red and gold. Line thickness indicates the strength/confidence of data support. Dotted lines mark the edges of the clusters. Among disconnected proteins, factors linked to actin organization are striped. The bait Tks5/SH3PXD2A is highlighted in bold. (C) Treemap representing the top biological processes represented among proteins biotinylated by Tks5-BirA*, as defined in (B), using PANTHER classification system for analysis and REVIGO for summarization and visualization. Each rectangle is a single cluster representative. The representatives are joined into ‘superclusters’ of loosely related terms, visualized with different colors. Size of the rectangles reflects the *p*-value of the GO term. (D) Proteins biotinylated by Tks5-BirA* (B) categorized as GO “Actin cytoskeleton organization” by PANTHER classification system.

To assess the biological relevance of this group of proteins, we surveyed the existence of previously described functional or physical protein-protein interactions (PPI) among them using the STRING platform. Such an analysis indicated that the identified Tks5 close neighbors form a significant functional network (PPI enrichment p-value =1.54e-09). The significance of this network was even greater when WASL/N-WASP and NCK1/Nck, that we validated as close neighbors directly by western blotting (Fig.2B), were included (PPI enrichment p-value =1.17e-14). Application of Markov Clustering logarithm (MCL) revealed that Tks5 close neighbors are organized in a main core cluster (Fig.3B - proteins in red and gold).

We next investigated molecular pathways associated with our biologically relevant network. Gene ontology analysis using PANTHER classification system indicated that “actin cytoskeleton organization” (GO: 0030036), critical for invadopodia regulation, was the most significantly overrepresented biological process in our network compared to its occurrence in the whole human genome (fold enrichment =11.91, P value =2.45e-10) strengthening the pertinence of our strategy (Fig.3C and D).

Furthermore, comparison of Tks5 close neighbors identified in the present study to GLAD4U literature mining protein listing obtained for “invadosome” query and manual literature searching revealed that 17.5% (7 out of 40) were previously linked to invadosomes (Fig.3A – red spots). Most of them have been implicated in invadosomes assembly through regulation of the actin cytoskeleton: Cortactin, a key component of invadopodia, which was among the most enriched biotinylated proteins with a fold enrichment (Log2) of 7.5 (Supplementary Table S1), is a regulator of Arp2/3 complex activity^27^; and the Cdc42 guanine exchange factor FGD1^28,29^, the formin binding protein FNBP1 (FBP17)^30–32^, the Ras-associated and pleckstrin homology containing protein 1 (RAPH1), also known as Lamellipodin (Lpd)^33^, and the Drebrin-like protein, DBNL (mAbp1)^34^ were all found to regulate invadosome formation through the control of factors implicated in elongation and branching of actin filaments. In contrast, the zonula occludens protein ZO-1/TJP1 was proposed to regulate degradative activity of invadopodia-like structures through its interaction with ADAM12 and MMP14^35,36^, whereas, the RNA binding protein IGF2BP2/IMP2 was shown to be involved in invadopodia formation through the control of the stability or localization of mRNAs of factors implicated in cell adhesion, mobility, invasion and ECM^37,38^. We also identified factors that have not been themselves involved in invadosome biology, but belong to protein families whose other members have (Fig.3A – orange spots). Such factors include the ADAM15 metalloproteinase^39,40^, the 5’-inositol lipid phosphatase Synaptojanin-1^41,42^, the CIN85/CMS family member CD2AP/CMS^43^ and the tight junction protein TJP2^35,36^. Of note, the majority of these proteins constitute the main core cluster of the integrative network identified after MCL clustering (Fig.3B). Additional proteins, including Reticulon-4 (RTN4)^44,45^, PDZ and LIM domain protein 4 (PDLIM4/RIL)^46,47^ and Dihydropyrimidase-like protein 3 (DPYSL3/CRMP4)^48^ were integrated to this main core cluster. Interestingly, these proteins, although not currently implicated in invadosome regulation, have been linked to actin cytoskeleton organization. Furthermore, as pointed out by the gene ontology analysis and manual literature searching, additional proteins involved in actin cytoskeleton organization were identified among Tks5 close neighbors (Fig.3B – striped circles and Fig.3D). These include the inhibitor of the Arp2/3 complex, Arpin^49^, the breast cancer anti-estrogen resistance protein 3 (BCAR3)^50,51^, the PDZ and LIM domain protein 1 (PDLIM1/ELFIN)^52,53^, the SHC-transforming protein 1 (SHC1) ^54^, the LIM and Calponin homology domain containing protein 1 (LIMCH1)^55^, the E3 ubiquitin ligase Mind Bomb 1 (MIB1)^56^, the microtubule-associated protein 4 (MAP4)^57^ and the anthrax toxin and collagen I receptor, ANTXR1/TEM8^58,59^. These observations strongly suggest that, through their control of actin organization, these proteins could be involved in invadosomes formation.

Thus, although other biological processes such as “D-ribose metabolism” or “tRNA wobble base modification”, were found significantly overrepresented by PANTHER classification system, the main pathway represented among Tks5 interactome is linked to actin cytoskeleton organization, which is critical for invadopodia regulation (Fig.3C). Furthermore, a large fraction (more than 50%) of Tks5 close neighbors identified here (Fig.3A and Supplementary Table S1) is associated with cancer cell invasion and metastasis.

### Invadopodia components show different modes of interaction with Tks5

To obtain further insights in the comprehension of invadopodia formation, we took advantage of the cellular model expressing the short form of Tks5, deleted of its PX domain (ΔPX-Tks5) (Fig.1). PX domains mediate interactions with specific membrane phosphoinositides. Tks5 PX domain was reported to preferentially interact with PI(3)P and PI(3,4)P2^22^. Such interaction contributes to Tks5 activation and thus to podosomes and invadopodia formation^13,21,60^. Our cellular model could therefore help us differentiate factors that associate with Tks5 before its recruitment to invadosome precursors, from those that associate with Tks5 at invadosome precursors. We therefore analyzed mass spectrometry data resulting from ΔPX-Tks5-BirA* pulldowns and identified 44 enriched proteins at 1% pFDR in ΔPX-Tks5-BirA* pulldown relatively to BirA* control pulldown (Supplementary Table S2). We then intersected this list of proteins with the *in situ* Tks5 interactome (Fig.4A and Supplementary Table S3). Twenty seven proteins were shared (Fig.4A - red spots and Supplementary Table S3), corresponding to more than half of the total proteins identified in each condition (67.5% and 61.3% of Tks5-BirA*- and ΔPX-Tks5-BirA*-associated proteins respectively). Some proteins were similarly enriched in the Tks5-BirA*- and ΔPX-Tks5-BirA*-associated groups, including the key invadopodia component Cortactin. Western blotting after pulldown of biotinylated proteins confirmed that Cortactin was biotinylated at similar levels by Tks5-BirA* and ΔPX-Tks5-BirA*, indicating that Cortactin is in the vicinity of Tks5 independently of its presence at invadopodia (Fig.4B). Other proteins were found exclusively or more strongly associated with Tks5-BirA* than ΔPX-Tks5-BirA*, among which thirteen were specifically biotinylated by Tks5-BirA*, but not by ΔPX-Tks5-BirA* nor BirA* (Fig.4A - blue spots and Supplementary Table S3). Reticulon-4, FGD1 and SHC1 were among the most highly enriched (Difference (Log2) >3.35). We confirmed by western blotting after pulldown of biotinylated proteins that RTN4 and FGD1 were specifically biotinylated by Tks5-BirA* (Fig.4B) and that MAP4 and CD2AP/CMS were biotinylated by both fusion proteins, but were more tightly associated with Tks5-BirA* (Fig.4B). We can speculate that the interaction of MAP4 and CD2AP is stabilized once Tks5 is recruited to PI(3,4)P2-enriched domains. Finally, for some of the shared proteins, fold enrichment was higher with ΔPX-Tks5-BirA* than with Tks5-BirA*, in accordance with the proposed mechanism whereby ΔPX-Tks5 might inhibit invadosome formation by mislocalizing actin-regulatory proteins^61^. Overall the comparison of Tks5α and Tks5β (ΔPX-Tks5) interactomes identifies three groups of Tks5 partners: (1) those interacting with Tks5 before its recruitment to PI(3,4)P2-rich domain, (2) those interacting weakly with Tks5 before its recruitment to PI(3,4)P2-rich domain, but whose interaction is stabilized once Tks5 is recruited, and (3) those interacting with Tks5 at invadopodia.

**Figure 4:**
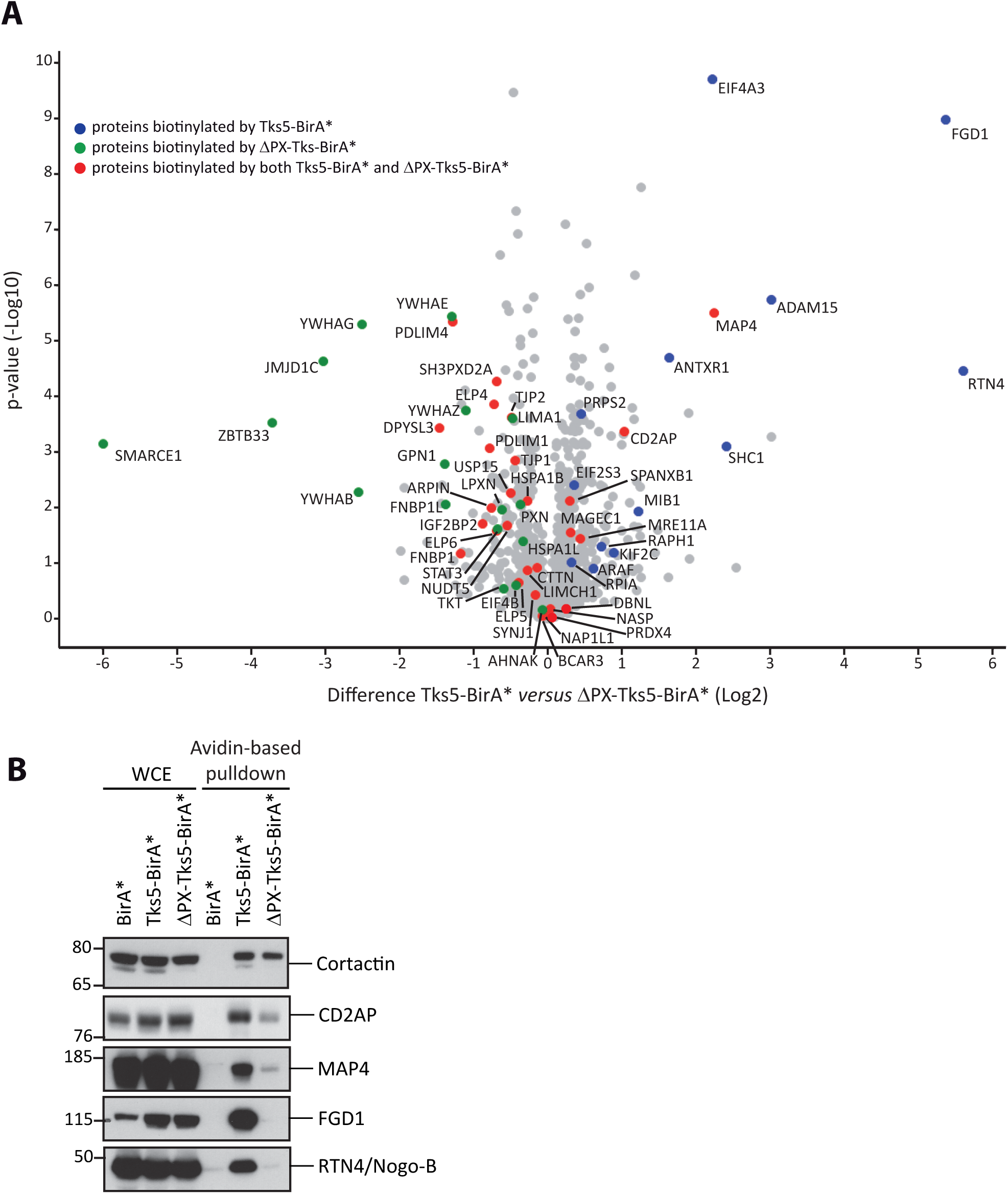
Invadopodia components show different modes of interaction with Tks5. MDA-MB-231 cells expressing Tks5-BirA*, ΔPX-Tks5-BirA* or BirA* were seeded on gelatin-coated plates and treated with biotin for 16 h. Biotinylated proteins were isolated by affinity capture and identified by mass spectrometry. Volcano plot showing differential level (Log2) and the p-value (-Log10), on x and y axis, respectively, for Tks5-BirA* *versus* ΔPX-Tks5-BirA*, without taking into account proteins identified as biotinylated in BirA* control condition. Data result from two different experiments processed three times. Proteins biotinylated by Tks5-BirA*, by ΔPX-Tks5-BirA* and by both Tks5-BirA* and ΔPX-Tks5-BirA*, relative to the BirA* control condition, are highlighted in blue, green and red, respectively. For more detail refer to Supplementary Table S3. (B) Verification of the biotinylation levels induced by Tks5-BirA* or ΔPX-Tks5-BirA*, of selected proteins identified in (A). The presence of proteins of interest was assessed in whole cell extract (WCE) or by avidin-based pulldown followed by western blotting with the indicated antibodies.

## DISCUSSION

Invadopodia, actin-rich protrusive structures developed at cancer cells surface in order to degrade the surrounding ECM^1–3^, are correlated to the metastatic potential of diverse cancer types, hence the importance of better understanding their biology^4,5^. In the current study, in order to gain insights into invadopodia molecular landscape, we determined the *in situ* interactome of Tks5, a key player in invadosome formation, by a global approach based on proximity-dependent labeling. Previous studies have identified Tks5 interacting partners^21,22,25,59^ using conventional approaches, based on identification of proteins co-precipitating with Tks5 or Tks5 domains, e.g. the SH3 domains. Proximity-dependent labeling methods in living cells are powerful technologies that allow to explore protein-protein interactions (PPI), or at least protein-protein close proximity, *in situ*, before cell lysis. Unlike more classical approaches, it allows to identify PPI in more physiologically relevant conditions, low affinity and/or transient PPI, as well as the proteome of membrane-less organelles and subcellular domains^10,11^. We choose to fuse Tks5 to the promiscuous biotin ligase BirA*. Fusion of BirA* to Tks5 did not affect its function: it was strongly enriched in invadopodia and promoted invadopodia formation and activity (Fig.1). Biotinylation is supposed to occur within an estimated range of 10nm^62^. Thus Tks5-BirA* should mostly biotinylate direct partners of Tks5, and proteins that belong to invadopodia, which have an estimated diameter of 0.05 to 1μm. We verified that Nck and N-WASP, invadopodia proteins and co-precipitation partners of Tks5, are indeed biotinylated by Tks5-BirA*, but only when present at invadopodia (Fig.2). This result also confirms that these interactions actually take place in intact cells.

Mass spectrometry analysis identified at least 40 proteins significantly associating with Tks5 (Fig.3 and Supplementary Table S1). A large number of the identified proteins are part of a functionally and physically-linked integrative network. Around twenty percent of Tks5 proximate proteins, or their homologs, have been previously linked to invadosomes and more than 50% are involved in cancer invasion and metastasis, revealing Tks5 *in situ* interactome as a source of potential new invadopodia components and regulators and strengthening the relevance of our approach. Very few studies have addressed the proteome of invadosomes, as their isolation and purification have been challenging. Two strategies have been used before: either cell fractionation by mechanical removal of the cell bodies in the case of invadopodia and podosomes^63,64^ or laser microdissection in the case of Src-induced rosettes ^65^. Although our list of proteins shares only few proteins with these studies (none with Attanasio et al., 1 with Cervero et al., 5 with Ezzoukhry et al.), a common feature between these studies and our results is the overrepresentation of factors involved in actin cytoskeleton organization (Fig.3). We could also identify proteins linked to metabolism and transcription/protein synthesis. However, in contrast to these studies in which these categories of proteins were highly enriched, they were poorly represented in our study. This difference might be linked to the strategy used. Indeed, whereas the other studies were based on the analysis of more or less pure cellular fractions, our strategy is limited to the identification of partners of a particular protein. Tks5 being usually depicted as localizing in the functional core of invadosomes in close proximity of factors regulating actin polymerization and ECM degradation, it is not surprising to predominantly identify factors linked to these processes among Tks5 close neighbors. Even though the biotinylation radius of BirA* is low, we cannot exclude that some of the proteins identified in Tks5 neighborhood, might be indirect partners. However, assessing molecular organization of Tks5 close neighbors using InterPro site, we observed that 17 out of 40, including the known invadopodia components FGD1, RAPH1 and FNBP1, contain either Pro-rich regions or SH3 domains, that could mediate a direct interaction with Tks5 (Supplementary Table S1). Further studies will be necessary to address their specific mode of interaction.

Assembly of invadopodia is a sequential process starting with the formation of an invadopodium precursor that will be stabilized and finally mature into a degradation-competent invadopodium^1,6,7^. Tks5 plays a major role in the stabilization step through interaction of its PX domain to PI(3,4)P2-enriched domains, and as a scaffold protein allowing the recruitment of factors necessary for invadopodium maturation^60^. Our comparison of the *in situ* interactomes of Tks5α (Tks5) and of its short form Tks5β (ΔPX-Tks5) suggests that, additionally to acting as a scaffolding factor in invadopodia-to-be PI(3,4)P2-rich domains, Tks5 associates with key invadopodia regulators, e.g. Cortactin, independently of its recruitment to invadopodia. Although these results allow us to get a first insight into the molecular mechanism of recruitment of proteins found in the vicinity of Tks5, further work would be necessary to address the spatiotemporal organization of invadopodia.

Overall, the present study demonstrates that proximity-dependent labeling methods in living cells are suitable to get an insight of invadopodia molecular composition. Indeed we identified known invadopodia components/regulators and new potential key invadopodia players for which validation of their implication in invadopodia activity and eventually in metastases formation, is needed. However, the fact that we did not identify Nck and N-WASP in the global strategy, while we defined them as Tks5 close neighbors, although at low levels, by a targeted strategy, points to limits in sensitivity of our strategy and suggest possible routes of improvements. Several options are under investigation to increase the sensitivity of MS detection, including the use of a more efficient promiscuous biotin ligase (e.g. TurboID) or the enhancement of the biotinylation range through the expansion of the linker region between the enzyme and the bait^66,67^. These strategies combined to the identification of the *in situ* protein neighbors of additional specific invadopodia components, such as Tks4, would permit to get a larger coverage of the invadopodia proteome. We can also envision to use such proximity-dependent labeling approaches to study the molecular composition of podosomes developed by normal cells and of the various types of invadopodia cancer cells differentially assemble during the invasion process depending on the matrix environment^68,69^. The slow kinetics of BirA* precludes its use for studying the dynamic molecular organization of invadopodia which have a lifetime in the order of one hour. Dynamics could be addressed by next generation proximity-dependent labeling techniques, such as the modified peroxidase APEX2 or TurboID, that might allow to resolve spatiotemporal regulation of invadopodia at the minute-scale^66,70^. The very existence of invadopodia or invadopodia-like structure in a physiological context, i.e. during tumor cell invasion, has been debated for years. As proximity-dependent approaches can also be applied *in vivo*^71–73^, our study opens the door to the development of proximity-dependent labeling approaches in *in vivo* models of invasion more relevant to metastatic progression.

## MATERIALS AND METHODS

### Cell lines

MDA-MB-231 cells were cultured in Dulbecco’s Modified Eagle Medium (ThermoFisher), supplemented with 10% fetal bovine serum (Eurobio, Courtaboeuf, France) at 37°C in a humidified atmosphere with 5% CO_2_ and checked regularly for mycoplasma contamination. Stable MDA-MB-231 cell lines were generated by transfection of the BirA* plasmid constructs by Lipofectamine 2000 (Invitrogen) and selection with 1 mg/ml geneticin. Individual clones were generated by limiting dilution and clones expressing moderate levels of the transgenes were selected for further analysis.

### Plasmid constructs

Wild type Tks5 and the mutant Tks5 lacking its PX domain, without stop codon, were amplified by PCR from peGFP-N1-Tks5 (a kind gift from S. Courtneidge) with appropriate primers (Supplementary Table S4), used to generate pDONR-Tks5 and pDONR-ΔPX-Tks5, respectively before their subcloning in the destination vector pcDNA3.1 MCS-BirA(R118G)-HA (modified from Addgene plasmid # 36047, a kind gift from K. J. Roux) by Gateway technology to obtain Tks5 and ΔPX-Tks5 fused to the N-terminus of BirA*-HA. All constructs were sequence verified.

### Antibodies

Rabbit antibodies against Tks5 (M-300), MAP4 (H-300), HA (Y-11) and mouse antibody against Grb2 (C-7) were purchased from Santa-Cruz Biotechnology. Mouse antibody against Cortactin (clone 4F11) was purchased from Millipore. Antibodies against N-WASP (mouse, ab56454), NogoA/B (rabbit, ab47085), CD2AP (rabbit, ab205017) were from Abcam. Mouse antibody against Nck (610099) was from BD Transduction Laboratories. Anti-α-Tubulin (mouse, clone DM1A) and anti-FGD1 (rabbit, HPA000911) were from Sigma Life Science. Biotinylated proteins were detected either by Alexa Fluor^®^594 Streptavidin (405240, Biolegend^®^) or Streptavidin-HRP (21130, Thermoscientific).

### Proximity biotinylation

Cells stably expressing BirA* fusion proteins were seeded on gelatin-coated plates and treated overnight with 50 μM biotin, then lysed in a buffer composed of 50 mM Tris pH 7.4, 500 mM NaCl, 0.4 % SDS, 2 % Triton X-100, 5 mM EDTA and 1 mM DTT supplemented with protease and phosphatase inhibitors. Biotinylated proteins were isolated by incubating the cell lysates with Avidin-coated beads (ThermoFisher) for 1h at 4°C. After three washes with lysis buffer and one wash with 50 mM Tris pH 7.4 plus 50 mM NaCl, beads were resuspended in SDS loading buffer supplemented with 2 mM D-biotin (Invitrogen).

### Western blotting

After protein extracts preparation, quantification and denaturation in SDS loading buffer, samples were run on Novex NuPAGE^®^ Bis-Tris 4-12% gels using a MOPS based running buffer. Proteins were transferred onto nitrocellulose membranes, incubated with primary antibodies and secondary antibodies coupled to HRP and detected by chemoluminescence.

### Mass spectrometry analysis

Biological samples were prepared in duplicate and each sample was further analyzed thrice by liquid chromatography (LC)-tandem mass spectrometry (MS/MS). Protein extracts were stacked on NuPAGE 4-12% Bis-Tris acrylamide gels (Life Technologies) and cut from the gel. Gels pieces were submitted to an in-gel trypsin digestion ^74^. Pooled extracts were dried down in a centrifugal vacuum system and samples were reconstituted and analyzed by LC-MS/MS using an LTQ Velos Orbitrap Mass Spectrometer (Thermo Electron, Bremen, Germany) online with an Ultimate 3000RSLCnano chromatography system (Thermo Fisher Scientific, Sunnyvale, CA). Peptides were separated on a Dionex Acclaim PepMap RSLC C18 column (15 cm × 75 µm I.D, 100 Å pore size, 2 µm particle size) at 300 nL/min flow rate. Peptides were eluted by a two steps linear gradient (4-22% acetonitrile/H_2_O; 0.1 % formic acid for 110 min and 22-32% acetonitrile/H_2_O; 0.1 % formic acid for 10 min). For peptide ionization in the EASY-Spray nanosource, spray voltage was set at 1.9 kV at 275 °C. MS spectra were acquired in the Orbitrap in the range of *m/z* 300-1700 at a FWHM resolution of 30 000 and using the 445.120025 ions as internal mass calibration. Collision-induced dissociation fragmentation was performed in the ion trap on the 10 most intense precursor ions. The signal threshold for an MS/MS event was set to 500 counts. Dynamic exclusion was enabled with a repeat count of 1, exclusion list size 500 and exclusion duration of 30 s.

### Protein identification and quantification

Relative intensity-based label-free quantification (LFQ) was processed using MaxQuant, version 1.6.3.4 ^75,76^ using mainly default parameters. The acquired raw LC-MS/MS data were first processed using the integrated Andromeda search engine ^77^. Spectra were searched against the Human database (UniProt Proteome reference, date 2019.01; 20412 entries) supplemented with a set of 245 frequently observed contaminants. The match between runs option was enabled to transfer identifications across different LC-MS/MS replicates based on their masses and retention time using default parameters. The quantification was performed using a minimum ratio count of 1 (unique+razor) and the second peptide option to allow identification of two co-fragmented co-eluting peptides with similar masses. For identification, the false discovery rate (FDR) at the peptide and protein levels were set to 1%. The statistical analysis was done with Perseus program (version 1.6.1.3) (www.maxquant.org) using LFQ normalised intensities. First, proteins marked as contaminant, reverse hits, and “only identified by site” were discarded. Protein LFQ normalized intensities were base 2 logarithmized to obtain a normal distribution. Quantifiable proteins were defined as those detected in at least 70% of the samples in one or more condition. Missing values were replaced using data imputation using default parameters. Differential proteins were selected using a two-sample t-test using permutation based FDR-controlled at 0.01 and employing 250 permutations. The p value was adjusted using a scaling factor s0 with a value of 1 ^78^.

### Proteome analysis

Gene ontology (GO) and enrichment analyses were performed using the Protein Analysis Through Evolutionary Relationships (PANTHER 14.1)(http://pantherdb.org/) ^81,82^ using the whole human genome as background. REVIGO web server was used to summarize and visualize GO terms (http://revigo.irb.hr/) ^83^. Interaction network were generated using STRING’s website (version 11.0) at high confidence (0.7) and network is clustered using a Markov Clustering algorithm "MCL inflation parameter = 3” (https://string-db.org/) ^84,85^.

### Matrix degradation assay

Acid-treated coverslips were incubated with 50 μg/ml poly-D-lysine 20 min at 37°C, then with 0.5% glutaraldehyde 15 min at RT and then inverted on a 40 μl drop of 0.2% gelatin plus Oregon Green 488–conjugated gelatin (Life technologies) (10:1) mixture 10 min at RT. The gelatin matrix was then quenched with 5 mg/ml sodium borohydride 5 min at RT and dehydrated in 70 % EtOH for storage. Thirty thousand MDA-MB-231 cells were plated for 4h on gelatin-coated coverslips rehydrated in complete growth medium for 1h before use. Cells were then fixed with 4% formaldehyde in PBS, permeabilized with 0.1 % Triton X-100 and blocked with 1 % BSA. Immunolabeling was performed with antibodies directed against target proteins and secondary antibodies labeled with DyLight^®^ 405 or AlexaFluor^®^ 594 (Jackson ImmunoResearch). Images were acquired on a Zeiss structured light ApoTome™ microscope equipped with a 63x 1.4 plan ApoChromat objective. Percentage of cells degrading was assessed by imaging 10 random fields per condition. Matrix degradation was analyzed by quantifying the mean degraded area in pixels and the number of degradation foci per cell using home-made Fiji macro. Precursor invadopodia are defined as Cortactin and Tks5 colocalization spots, whereas mature invadopodia correspond to spots of cortactin and Tks5 colocalizing with degradation foci. 25 cells per condition per experiment were analyzed to quantify matrix degradation and invadopodia.

### Statistical analysis

All statistical analyses were performed using GraphPad Prism software. The unpaired one-tailed *t* test, with Welch correction, and the Mann Whitney test, were used to determine significant differences between data groups. Graphs were plotted using Prism, to show the mean and SEM. P values are indicated on the graph as * P<0.05, ** P<0.01, *** P<0.001.

## ACKNOWLEDGEMENTS

We thank D. Isnardon and M. Rodriguez (CRCM Microscopy Platform) for support and S.A. Courtneidge and K.J. Roux for sharing reagents. This work was supported by Canceropôle Provence-Alpes-Côte d’Azur (PACA), Institut National du Cancer, Conseil Régional PACA and Site de Recherche Intégrée sur le Cancer (SIRIC). Proteomic analyses were performed at the mass spectrometry facility of Marseille Proteomics supported by IBISA (Infrastructures Biologie Santé et Agronomie), Plateforme Technologique Aix-Marseille, Canceropôle PACA, Région Sud Provence-Alpes-Côte d’Azur, Fonds Européen de Développement Régional (FEDER) and Plan Cancer.

## AUTHOR CONTRIBUTIONS

S.T. and A.B. designed research; S.T., C.M., J.S., K.O., B.C., D.S. and E.B. performed research; S.T., C.M., J.S., K.O., B.C., S.A., L.C. and A.B. analyzed data; S.T. and A.B. wrote the paper.

