## Supplementary Information for "A proximity-labeling proteomic approach to investigate invadopodia molecular landscape in breast cancer cells"

**Supplementary Table S1: List of proteins biotinylated by Tks5-BirA* fusion protein**. Protein list results from the comparison of proteins biotinylated by Tks5-BirA* fusion protein with proteins biotinylated by BirA* control protein, after their isolation by affinity capture and identification by mass spectrometry. Proteins listed here correspond to proteins above the full line indicative of protein hits obtained at a pFDR of 1% on Fig.3A volcano plot. For each protein their link to invadosome, their implication in actin cytoskeleton organization and their link to cancer invasion and metastasis have been analyzed, by comparison to the list of proteins obtained for "invadosome" query by GLAD4U literature mining and manual literature searching. The presence of Proline-rich regions and SH3 domains determined by InterPro site is also referenced. Data result from two different experiments processed three times.

**Supplementary Table S2: List of proteins biotinylated by ∆PX-Tks5-BirA* fusion protein.** Protein list results from the comparison of proteins biotinylated by ∆PX-Tks5-BirA* fusion protein with proteins biotinylated by BirA* control protein, after their isolation by affinity capture and identification by mass spectrometry. Proteins listed here correspond to proteins hits obtained at a pFDR of 1%. Data result from two different experiments processed three times.

**Supplementary Table S3: Tables listing proteins biotinylated by both Tks5-BirA* and ∆PX-Tks5-BirA* (left), only by Tks5-BirA* (middle) and only by ∆PX-Tks5-BirA* (right) fusion proteins.** Proteins listed here correspond to proteins highlighted in red, blue and green, respectively, on Fig.4A volcano plot. Tables result from the intersection of Tks5-BirA* (Supplementary Table S1) and ∆PX-Tks5-BirA* (Supplementary Table S2) close neighbors. Proteins biotinylated by both Tks5-BirA* fusion proteins correspond to proteins identified as different (pFDR of 1%) in each of the fusion protein relative to BirA* control condition. Proteins designed as Tks5-BirA* or ∆PX-Tks5-BirA* only correspond to proteins different (pFDR of 1%) only in the fusion protein addressed relative to BirA* control condition. For each protein, its difference of abundance in Tks5-BirA* fusion proteins *versus* BirA* conditions is indicated and its difference of abundance in Tks5-BirA* *versus* ∆PX-Tks5-BirA* conditions has been calculated and is reported in the third column. [(1)] Grey-colored columns correspond to proteins with no significant differential of abundance in the Tks5-BirA* fusion protein analyzed relative to BirA* control condition.

**Supplementary Table S4:** Primers pairs used for amplifying Tks5 and ∆PX-Tks5 sequences.

**Supplementary Table S1: List of proteins biotinylated by Tks5-BirA* fusion protein.**

Protein list results from the comparison of proteins biotinylated by Tks5-BirA* fusion protein with proteins biotinylated by BirA* control protein, after their isolation by affinity capture and identification by mass spectrometry.
Proteins listed here correspond to proteins above the full line indicative of protein hits obtained at a pFDR of 1% on Fig.3A volcano plot.
For each protein their link to invadosome, their implication in actin cytoskeleton organization and their link to cancer invasion and metastasis have been analyzed, by comparison to the list of proteins obtained for "invadosome" query by GLAD4U literature mining and manual literature searching.
The presence of Proline-rich regions and SH3 domains determined by InterPro site is also referenced.
Data result from two different experiments processed three times.

| Protein IDs | Protein names | Gene names | Peptides | Sequence coverage [%] | Mol. weight [kDa] | p-value (-Log10) | Differential (Log2) Tks5-BirA* *versus* BirA* | Linked to invadosomes | Implicated in actin cytoskeleton reorganization | Linked to cancer invasion/metastasis | SH3 containing proteins | Proline-rich region containing proteins | References (PMID:) |
|---|---|---|---|---|---|---|---|---|---|---|---|---|---|
| Q5TCZ1 | SH3 and PX domain-containing protein 2A | SH3PXD2A | 120 | 81 | 125.29 | 13.91414333 | 16.10223103 | Yes | Yes | Yes | Yes | Yes | |
| P49959 | Double-strand break repair protein MRE11A | MRE11A | 18 | 35.9 | 80.592 | 7.93361717 | 7.613502502 | | | Yes | | | 22914783; 30176843 |
| Q14247 | Src substrate cortactin | CTTN | 28 | 50.5 | 61.585 | 10.4023223 | 7.51793925 | Yes | Yes | Yes | Yes | | 10442635; 29172953 |
| Q9NQC3 | Reticulon-4 | RTN4 | 7 | 8.6 | 129.93 | 7.876353516 | 7.337134679 | | Yes | Yes | | Yes | 28628795; 23042479; 27132996 |
| Q9Y5K6 | CD2-associated protein | CD2AP | 13 | 24.3 | 71.45 | 8.983150317 | 7.225888411 | (Yes) | Yes | | Yes | Yes | 10339567; 23090967 |
| O60732 | Melanoma-associated antigen C1 | MAGEC1 | 6 | 8.5 | 123.64 | 5.386851732 | 5.631188869 | | | Yes | | | 21738656; 17607369 |
| P98174 | FYVE, RhoGEF and PH domain-containing protein 1 | FGD1 | 7 | 12.3 | 106.56 | 8.73033328 | 4.930959066 | Yes | Yes | Yes | | Yes | 19141649; 22854039 |
| P27816 | Microtubule-associated protein 4 | MAP4 | 16 | 20.4 | 121 | 8.687608396 | 4.929625193 | | Yes | Yes | | Yes | 21937675;24140250; 26876215 |
| Q96RU3 | Formin-binding protein 1 | FNBP1 | 6 | 12.5 | 71.306 | 4.260582509 | 4.438922723 | Yes | Yes | Yes | Yes | Yes | 29651632; 21421245 |
| P29353 | SHC-transforming protein 1 | SHC1 | 4 | 16.8 | 62.821 | 6.160669832 | 3.875651677 | | Yes | Yes | | Yes | 24078705; 18604176; 23277357 |
| Q14195 | Dihydropyrimidinase-related protein 3 | DPYSL3 | 12 | 30.4 | 61.963 | 3.494760306 | 3.636353652 | | Yes | Yes (+ or -) | | | 16181627; 26934554; 22805864; 20543870 |
| Q9Y6M1 | Insulin-like growth factor 2 mRNA-binding protein 2 | IGF2BP2 | 3 | 8.5 | 66.121 | 4.416141395 | 2.985394319 | Yes | | Yes | | | 29217458; 16541107; 23069990 |
| Q9NS25 | Sperm protein associated with the nucleus on the X chromosome B/F | SPANXB1 | 2 | 27.2 | 11.84 | 6.075416422 | 2.941437085 | | | Yes (+ or -) | | | 26895102; 31406142; 27323831 |
| Q9NS26 | Sperm protein associated with the nucleus on the X chromosome A | SPANXA1 | | | | | | | | | | | |
| Q86YT6 | E3 ubiquitin-protein ligase MIB1 | MIB1 | 2 | 3.4 | 110.13 | 3.916397356 | 2.915704568 | | Yes | | | | 29078376 |
| Q8TE02 | Elongator complex protein 5 | ELP5 | 5 | 24.1 | 34.841 | 3.2596053 | 2.9046731 | | | Yes | | | 22854966 |
| Q9UKK9 | ADP-sugar pyrophosphatase | NUDT5 | 4 | 23.3 | 24.327 | 3.925240339 | 2.815107028 | | | | | | |
| Q96EB1 | Elongator complex protein 4 | ELP4 | 6 | 24.1 | 46.587 | 7.643664097 | 2.812804858 | | | Yes | | | 29805303 |
| Q9H6X2 | Anthrax toxin receptor 1 | ANTXR1 | 4 | 11.5 | 62.788 | 3.883642755 | 2.784154256 | | Yes | Yes | | Yes | 19817382; 23832666 |
| O00151 | PDZ and LIM domain protein 1 | PDLIM1 | 4 | 27.7 | 36.071 | 3.126147395 | 2.704363346 | | Yes | Yes (+ or -) | | | 11110697; 31509262; 26701804; 24662836 |
| Q13444 | Disintegrin and metalloproteinase domain-containing protein 15 | ADAM15 | 3 | 5.6 | 92.958 | 4.679859557 | 2.634385427 | (Yes) | | Yes (+ or -) | | Yes | 26930657; 18281484; 20851104 |
| Q7Z6K5 | Arpin | ARPIN | 1 | 5.3 | 24.943 | 6.153299742 | 2.345861912 | | Yes | Yes (-) | | | 24132237; 30416079; 26867158 |
| Q70E73 | Ras-associated and pleckstrin homology domains-containing protein 1 | RAPH1 | 3 | 4.7 | 135.25 | 2.33985344 | 2.165382385 | Yes | Yes | Yes | | Yes | 26996666; 30056565 |
| Q9UJU6 | Drebrin-like protein | DBNL | 1 | 2.6 | 48.207 | 3.83907753 | 2.131621997 | Yes | Yes | Yes (+ or -) | Yes | | 27129278; 22303001; 24912570 |
| Q07157 | Tight junction protein ZO-1 | TJP1 | 24 | 20.7 | 195.46 | 7.2425774 | 2.101765315 | Yes | Yes | Yes ? | Yes | Yes | 29765546; 20930113 |
| P55209 | Nucleosome assembly protein 1-like 1 | NAP1L1 | 2 | 5.4 | 45.374 | 3.573022604 | 2.077949206 | | | Yes | | | 31516385 |
| Q99733 | Nucleosome assembly protein 1-like 4 | NAP1L4 | | | | | | | | | | | |
| O43426 | Synaptojanin-1 | SYNJ1 | 2 | 3.1 | 173.1 | 4.141125533 | 2.071942647 | (Yes) | | | | Yes | 25605973; 9199318 |
| Q99661 | Kinesin-like protein KIF2C | KIF2C | 2 | 5 | 81.312 | 2.094197001 | 1.995908578 | | | Yes | | | 18506187; 17653072; 31073307 |
| Q13162 | Peroxiredoxin-4 | PRDX4 | 2 | 8.1 | 30.54 | 2.563453061 | 1.873274485 | | | Yes | | | 31311441; 21487000; 25054044 |
| O75815 | Breast cancer anti-estrogen resistance protein 3 | BCAR3 | 2 | 3.3 | 92.565 | 2.254819159 | 1.872347673 | | Yes | Yes (+/-) | | | 25499443; 17616674; 23762409 |
| P50479 | PDZ and LIM domain protein 4 | PDLIM4 | 2 | 10.6 | 35.398 | 2.950149833 | 1.868245602 | | Yes | Yes (-) | | | 14729062; 31376371; 20642860 |
| P10398 | Serine/threonine-protein kinase A-Raf | ARAF | 1 | 3 | 67.585 | 2.441532048 | 1.833749294 | | | | | | |
| Q0PNE2 | Elongator complex protein 6 | ELP6 | 2 | 9.8 | 29.793 | 2.127950462 | 1.777745883 | | | Yes | | | 22854966 |
| P11908 | Ribose-phosphate pyrophosphokinase 2 | PRPS2 | 8 | 36.2 | 34.769 | 5.812583924 | 1.750796954 | | | Yes | | | 30908912 |
| Q9Y4E8 | Ubiquitin carboxyl-terminal hydrolase 15 | USP15 | 1 | 2.1 | 112.42 | 2.472299838 | 1.748913447 | | | | | | |
| P38919 | Eukaryotic initiation factor 4A-III;Eukaryotic initiation factor 4A-III, N-terminally processed | EIF4A3 | 13 | 29.7 | 46.871 | 7.984805868 | 1.744141579 | | | | | | |
| Q9UPQ0 | LIM and calponin homology domains-containing protein 1 | LIMCH1 | 2 | 2.7 | 121.87 | 3.924842063 | 1.579572042 | | Yes | | | | 28228547 |
| P41091 | Eukaryotic translation initiation factor 2 subunit 3 | EIF2S3 | 9 | 30.3 | 51.109 | 7.34536761 | 1.521572113 | | | | | | |
| Q2VIR3 | Putative eukaryotic translation initiation factor 2 subunit 3-like protein | EIF2S3L | | | | | | | | | | | |
| P49247 | Ribose-5-phosphate isomerase | RPIA | 5 | 22.5 | 33.269 | 6.120177273 | 1.411331177 | | | Yes | | | 25429733 |
| Q9UDY2 | Tight junction protein ZO-2 | TJP2 | 14 | 17.2 | 133.96 | 7.114464632 | 1.4007864 | (Yes) | Yes | Yes (-) | Yes | | 23605953 |
| P49321 | Nuclear autoantigenic sperm protein | NASP | 4 | 11.4 | 85.237 | 6.253539977 | 1.394634247 | | | | | | |
| P0DMV9 | Heat shock 70 kDa protein 1B | HSPA1B | 37 | 70.2 | 70.051 | 9.848124347 | 1.286704063 | | | Yes | | | 19914824; 20564126; 25213699 |
| P0DMV8 | Heat shock 70 kDa protein 1A | HSPA1A | | | | | | | | | | | |

**Supplementary Table S2: List of proteins biotinylated by ∆PX-Tks5-BirA* fusion protein.**

Protein list results from the comparison of proteins biotinylated by ∆PX-Tks5-BirA* fusion protein with proteins biotinylated by BirA* control protein, after their isolation by affinity capture and identification by mass spectrometry.
Proteins listed here correspond to proteins hits obtained at a pFDR of 1%. Data result from two different experiments processed three times.

| Protein IDs | Protein names | Gene names | Peptides | Sequence coverage [%] | Mol. weight [kDa] | p-value (-Log10) | Differential (Log2) ∆PX-Tks5-BirA* *versus* BirA* |
|---|---|---|---|---|---|---|---|
| Q5TCZ1;A1X283 | SH3 and PX domain-containing protein 2A | SH3PXD2A | 120 | 81 | 125.29 | 14.00469377 | 16.79133638 |
| Q14247 | Src substrate cortactin | CTTN | 28 | 50.5 | 61.585 | 10.5672796 | 7.658259074 |
| P49959 | Double-strand break repair protein MRE11A | MRE11A | 18 | 35.9 | 80.592 | 7.46094941 | 7.174569766 |
| Q9Y5K6 | CD2-associated protein | CD2AP | 13 | 24.3 | 71.45 | 7.966618016 | 6.196915785 |
| Q96RU3 | Formin-binding protein 1 | FNBP1 | 6 | 12.5 | 71.306 | 4.670416997 | 5.608518759 |
| O60732 | Melanoma-associated antigen C1 | MAGEC1 | 6 | 8.5 | 123.64 | 5.166906572 | 5.325631301 |
| Q14195 | Dihydropyrimidinase-related protein 3 | DPYSL3 | 12 | 30.4 | 61.963 | 4.583939755 | 5.090364933 |
| Q86T24 | Transcriptional regulator Kaiso | ZBTB33 | 3 | 8.8 | 74.484 | 7.226418622 | 4.578668594 |
| Q969G3 | SWI/SNF-related matrix-associated actin-dependent regulator of chromatin subfamily E member 1 | SMARCE1 | 2 | 8 | 46.649 | 2.205633135 | 4.180418491 |
| Q15652 | Probable JmjC domain-containing histone demethylation protein 2C | JMJD1C | 3 | 1.8 | 284.52 | 5.743897208 | 4.147589048 |
| Q9Y6M1 | Insulin-like growth factor 2 mRNA-binding protein 2 | IGF2BP2 | 3 | 8.5 | 66.121 | 5.538307203 | 3.866715272 |
| Q96EB1 | Elongator complex protein 4 | ELP4 | 6 | 24.1 | 46.587 | 8.216873652 | 3.530961037 |
| O00151 | PDZ and LIM domain protein 1 | PDLIM1 | 4 | 27.7 | 36.071 | 3.974932048 | 3.492241383 |
| Q9UKK9 | ADP-sugar pyrophosphatase | NUDT5 | 4 | 23.3 | 24.327 | 4.395425637 | 3.363261541 |
| Q8TE02 | Elongator complex protein 5 | ELP5 | 5 | 24.1 | 34.841 | 3.549250869 | 3.296580474 |
| P50479 | PDZ and LIM domain protein 4 | PDLIM4 | 2 | 10.6 | 35.398 | 4.918884984 | 3.1442132 |
| Q7Z6K5 | Arpin | ARPIN | 1 | 5.3 | 24.943 | 5.78745232 | 3.105995337 |
| P31946 | 14-3-3 protein beta/alpha;14-3-3 protein beta/alpha, N-terminally processed | YWHAB | 6 | 40.7 | 28.082 | 3.102797521 | 2.770226796 |
| Q5T0N5 | Formin-binding protein 1-like | FNBP1L | 3 | 8.6 | 70.065 | 3.163598993 | 2.732745806 |
| P61981 | 14-3-3 protein gamma;14-3-3 protein gamma, N-terminally processed | YWHAG | 5 | 32.4 | 28.302 | 3.841400861 | 2.722501755 |
| P27816 | Microtubule-associated protein 4 | MAP4 | 16 | 20.4 | 121 | 6.926183764 | 2.682752927 |
| Q9NS25 | Sperm protein associated with the nucleus on the X chromosome B/F | SPANXB1 | 2 | 27.2 | 11.84 | 5.73212081 | 2.642450651 |
| Q9NS26 | Sperm protein associated with the nucleus on the X chromosome A | SPANXA1 | | | | | |
| Q07157 | Tight junction protein ZO-1 | TJP1 | 24 | 20.7 | 195.46 | 8.285232815 | 2.545808792 |
| Q0PNE2 | Elongator complex protein 6 | ELP6 | 2 | 9.8 | 29.793 | 2.754979228 | 2.469645182 |
| Q9Y4E8 | Ubiquitin carboxyl-terminal hydrolase 15 | USP15 | 1 | 2.1 | 112.42 | 3.096935139 | 2.253142675 |
| O43426 | Synaptojanin-1 | SYNJ1 | 2 | 3.1 | 173.1 | 4.567370702 | 2.237295151 |
| P29401 | Transketolase | TKT | 4 | 11.1 | 67.877 | 2.172138733 | 2.136732419 |
| P55209 | Nucleosome assembly protein 1-like 1 | NAP1L1 | 2 | 5.4 | 45.374 | 3.424767194 | 2.123708725 |
| Q99733 | Nucleosome assembly protein 1-like 4 | NAP1L4 | | | | | |
| Q9HCN4 | GPN-loop GTPase 1 | GPN1 | 1 | 4.3 | 41.74 | 3.390386248 | 2.03527228 |
| P23588 | Eukaryotic translation initiation factor 4B | EIF4B | 2 | 6.2 | 69.15 | 2.576545034 | 1.942392349 |
| O75815 | Breast cancer anti-estrogen resistance protein 3 | BCAR3 | 2 | 3.3 | 92.565 | 2.435325241 | 1.941552321 |
| P62258 | 14-3-3 protein epsilon | YWHAE | 9 | 37.6 | 29.174 | 7.019493727 | 1.934883118 |
| Q9UJU6 | Drebrin-like protein | DBNL | 1 | 2.6 | 48.207 | 1.841725873 | 1.888232867 |
| Q9UDY2 | Tight junction protein ZO-2 | TJP2 | 14 | 17.2 | 133.96 | 8.786069397 | 1.886053403 |
| Q9UPQ0 | LIM and calponin homology domains-containing protein 1 | LIMCH1 | 2 | 2.7 | 121.87 | 4.156091474 | 1.855589549 |
| P40763 | Signal transducer and activator of transcription 3 | STAT3 | 1 | 3.4 | 88.067 | 2.973856904 | 1.832819303 |
| Q13162 | Peroxiredoxin-4 | PRDX4 | 2 | 8.1 | 30.54 | 2.654100435 | 1.831843694 |
| Q9UHB6 | LIM domain and actin-binding protein 1 | LIMA1 | 4 | 7.4 | 85.225 | 3.11598462 | 1.791516304 |
| P0DMV9 | Heat shock 70 kDa protein 1B | HSPA1B | 37 | 70.2 | 70.051 | 8.983589311 | 1.561729113 |
| P0DMV8 | Heat shock 70 kDa protein 1A | HSPA1A | | | | | |
| P49023 | Paxillin | PXN | 8 | 23.2 | 64.505 | 5.453557229 | 1.528698921 |
| P63104 | 14-3-3 protein zeta/delta | YWHAZ | 10 | 45.3 | 27.745 | 4.353593263 | 1.456914902 |
| P34931 | Heat shock 70 kDa protein 1-like | HSPA1L | 15 | 29.8 | 70.374 | 4.714809086 | 1.392109235 |
| O60711 | Leupaxin | LPXN | 1 | 5.7 | 43.332 | 4.515592353 | 1.385031064 |
| P49321 | Nuclear autoantigenic sperm protein | NASP | 4 | 11.4 | 85.237 | 6.546653084 | 1.354332606 |
| Q09666 | Neuroblast differentiation-associated protein AHNAK | AHNAK | 104 | 39.7 | 629.09 | 6.490538917 | 1.249121666 |

**Supplementary Table S3: Tables listing proteins biotinylated by both Tks5-BirA* and ∆PX-Tks5-BirA*(left), only by Tks5-BirA* (middle) and only by ∆PX-Tks5-BirA* (right) fusion proteins.**

Proteins listed here correspond to proteins highlighted in red, blue and green, respectively, on Fig.4A volcano plot
Tables result from the intersection of Tks5-BirA* (Supplementary Table S1) and ∆PX-Tks5-BirA* (Supplementary Table S2) close neighbors.
Proteins biotinylated by both Tks5-BirA* fusion proteins correspond to proteins identified as different (pFDR < 1%) in each of the fusion protein relative to BirA* control condition.
Proteins designed as Tks5-BirA* or DPX-Tks5-BirA* only correspond to proteins different (pFDR < 1%) only in the fusion protein addressed relative to BirA* control condition.
For each protein, its difference of abundance in Tks5-BirA* fusion proteins versus BirA* conditions is indicated, and its difference of abundance in Tks5-BirA* versus DPX-Tks5-BirA* conditions has been calclutated and is reported in the third column.
(1) Grey-colored columns correspond to proteins with no significant differential of abundance in the Tks5-BirA* fusion protein analyzed relative to BirA* control condition.

| In both (27) | Differential (Log2) Tks5-BirA* *versus* BirA* | Differential (Log2) ∆PX-Tks5-BirA* *versus* BirA* | Differential (Log2) Tks5-BirA* *versus* ∆PX-Tks5-BirA* |
|---|---|---|---|
| MAP4 | 4.93 | 2.68 | 2.25 |
| CD2AP | 7.23 | 6.20 | 1.03 |
| MRE11A | 7.61 | 7.17 | 0.44 |
| MAGEC1 | 5.63 | 5.33 | 0.31 |
| SPANXA1/SPANXB1 | 2.94 | 2.64 | 0.30 |
| DBNL | 2.13 | 1.89 | 0.24 |
| PRDX4 | 1.87 | 1.83 | 0.04 |
| NASP | 1.39 | 1.35 | 0.04 |
| NAP1L1/NAP1L4 | 2.08 | 2.12 | -0.05 |
| BCAR3 | 1.87 | 1.94 | -0.07 |
| CTTN | 7.52 | 7.66 | -0.14 |
| SYNJ1 | 2.07 | 2.24 | -0.17 |
| HSPA1A/HSPA1B | 1.29 | 1.56 | -0.28 |
| LIMCH1 | 1.58 | 1.86 | -0.28 |
| ELP5 | 2.90 | 3.30 | -0.39 |
| TJP1 | 2.10 | 2.55 | -0.44 |
| TJP2 | 1.40 | 1.89 | -0.49 |
| USP15 | 1.75 | 2.25 | -0.50 |
| NUDT5 | 2.82 | 3.36 | -0.55 |
| ELP6 | 1.78 | 2.47 | -0.69 |
| ELP4 | 2.81 | 3.53 | -0.72 |
| ARPIN | 2.35 | 3.11 | -0.76 |
| PDLIM1 | 2.70 | 3.49 | -0.79 |
| IGF2BP2 | 2.99 | 3.87 | -0.88 |
| FNBP1 | 4.44 | 5.61 | -1.17 |
| PDLIM4 | 1.87 | 3.14 | -1.28 |
| DPYSL3 | 3.64 | 5.09 | -1.45 |

| In Tks5-BirA* only (13) | Differential (Log2) Tks5-BirA* *versus* BirA* | Differential (Log2) ∆PX-Tks5-BirA* *versus* BirA* (1) | Differential (Log2) Tks5-BirA* *versus* ∆PX-Tks5-BirA* |
|---|---|---|---|
| RTN4 | 7.34 | 1.74 | 5.60 |
| FGD1 | 4.93 | 0.00 | 4.93 |
| SHC1 | 3.88 | 1.47 | 2.40 |
| MIB1 | 2.92 | 2.64 | 0.27 |
| ANTXR1 | 2.78 | 1.15 | 1.64 |
| ADAM15 | 2.63 | -0.38 | 3.01 |
| RAPH1 | 2.17 | 1.44 | 0.72 |
| KIF2C | 2.00 | 1.11 | 0.88 |
| ARAF | 1.83 | 1.22 | 0.61 |
| PRPS2 | 1.75 | 1.30 | 0.45 |
| EIF4A3 | 1.74 | -0.47 | 2.21 |
| EIF2S3/EIF2S3L | 1.52 | 1.16 | 0.36 |
| RPIA | 1.41 | 1.10 | 0.31 |

| In ∆PX-Tks5-BirA* only (17) | Differential (Log2) Tks5-BirA* *versus* BirA* (1) | Differential (Log2) ∆PX-Tks5–BirA* *versus* BirA* | Differential (Log2) Tks5-BirA* *versus* ∆PX-Tks5-BirA* |
|---|---|---|---|
| ZBTB33 | 0.86 | 4.58 | -3.72 |
| SMARCE1 | 0.00 | 4.18 | -4.18 |
| JMJD1C | 1.13 | 4.15 | -3.02 |
| YWHAB | 0.22 | 2.77 | -2.55 |
| FNBP1L | 1.36 | 2.73 | -1.38 |
| YWHAG | 0.22 | 2.72 | -2.50 |
| TKT | 1.54 | 2.14 | -0.59 |
| GPN1 | 0.64 | 2.04 | -1.39 |
| EIF4B | 1.51 | 1.94 | -0.43 |
| YWHAE | 0.64 | 1.93 | -1.29 |
| STAT3 | 1.15 | 1.83 | -0.68 |
| LIMA1 | 1.31 | 1.79 | -0.48 |
| PXN | 1.16 | 1.53 | -0.37 |
| YWHAZ | 0.35 | 1.46 | -1.11 |
| HSPA1L | 1.06 | 1.39 | -0.33 |
| LPXN | 0.77 | 1.39 | -0.62 |
| AHNAK | 1.18 | 1.25 | -0.07 |

**Supplementary Table S4 : Primer pairs used for amplifying Tks5 and ∆PX-Tks5 sequences**

| Name | Sequence |
|---|---|
| attB1-Nter Tks5 | 5’-GGGGACAAGTTTGTACAAAAAAGCAGGCTTCACCATGCTCGC CTACTGCGTGCAGGATG-3’ |
| attB1-delPX Tks5 | 5’-GGGGACAAGTTTGTACAAAAAAGCAGGCTTCACCATGGAGGC TCGACCCGAGGATGTC-3’ |
| attB2-Cter Tks5 | 5’-GGGGACCACTTTGTACAAGAAAGCTGGGTCGTTCTTTTTCTCAAGGTAGTTGGAAG-3’ |

Figure 1C

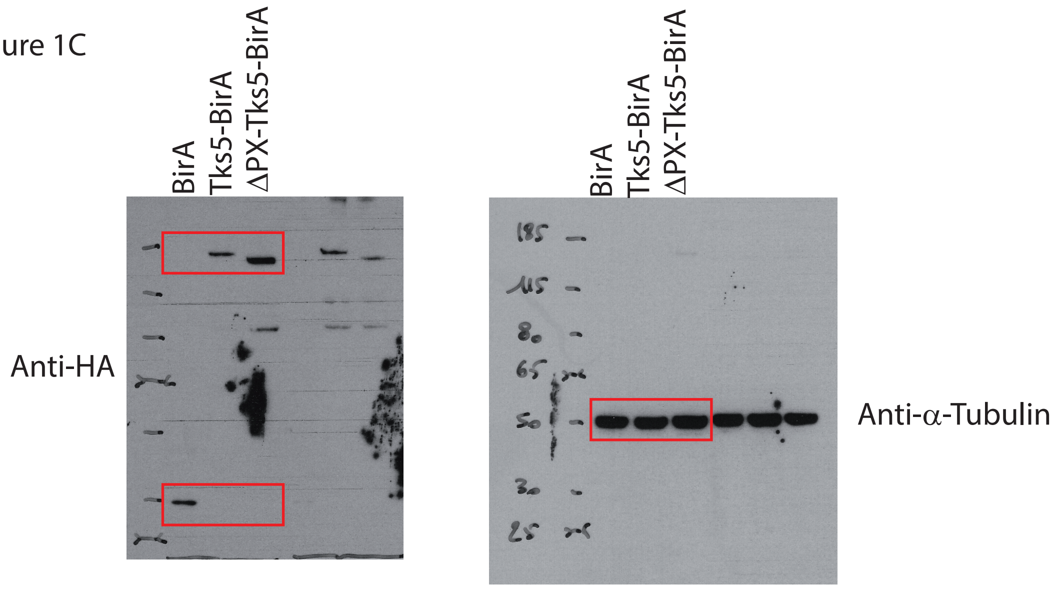


Figure 2B

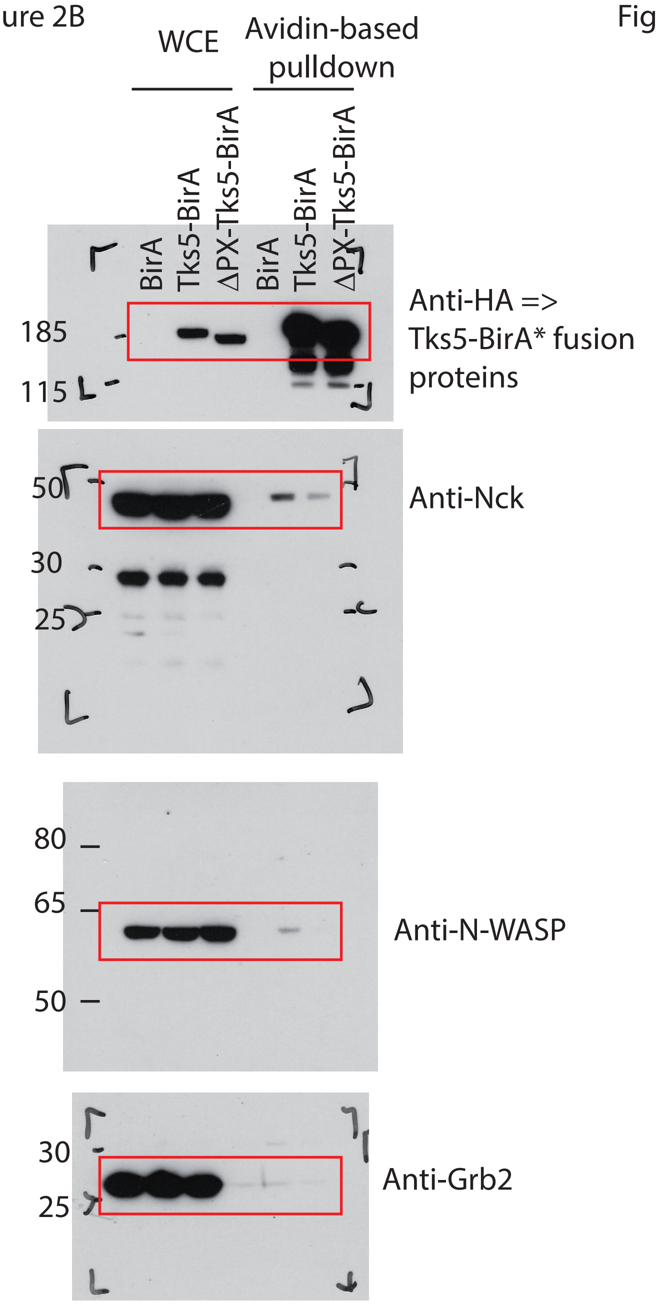


Figure 4B

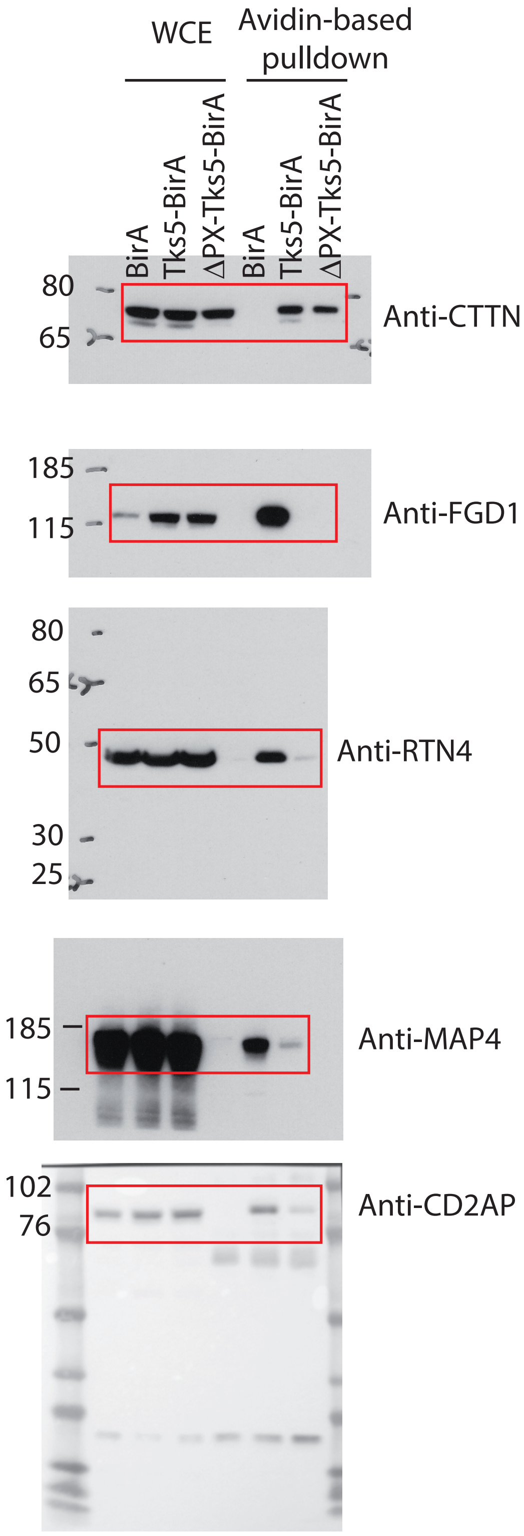
